## Supplementary figures and images for "RBM7 deficiency promotes breast cancer metastasis by coordinating MFGE8 splicing switch and NF-kB pathway"

### Figure 1-figure supplement 1

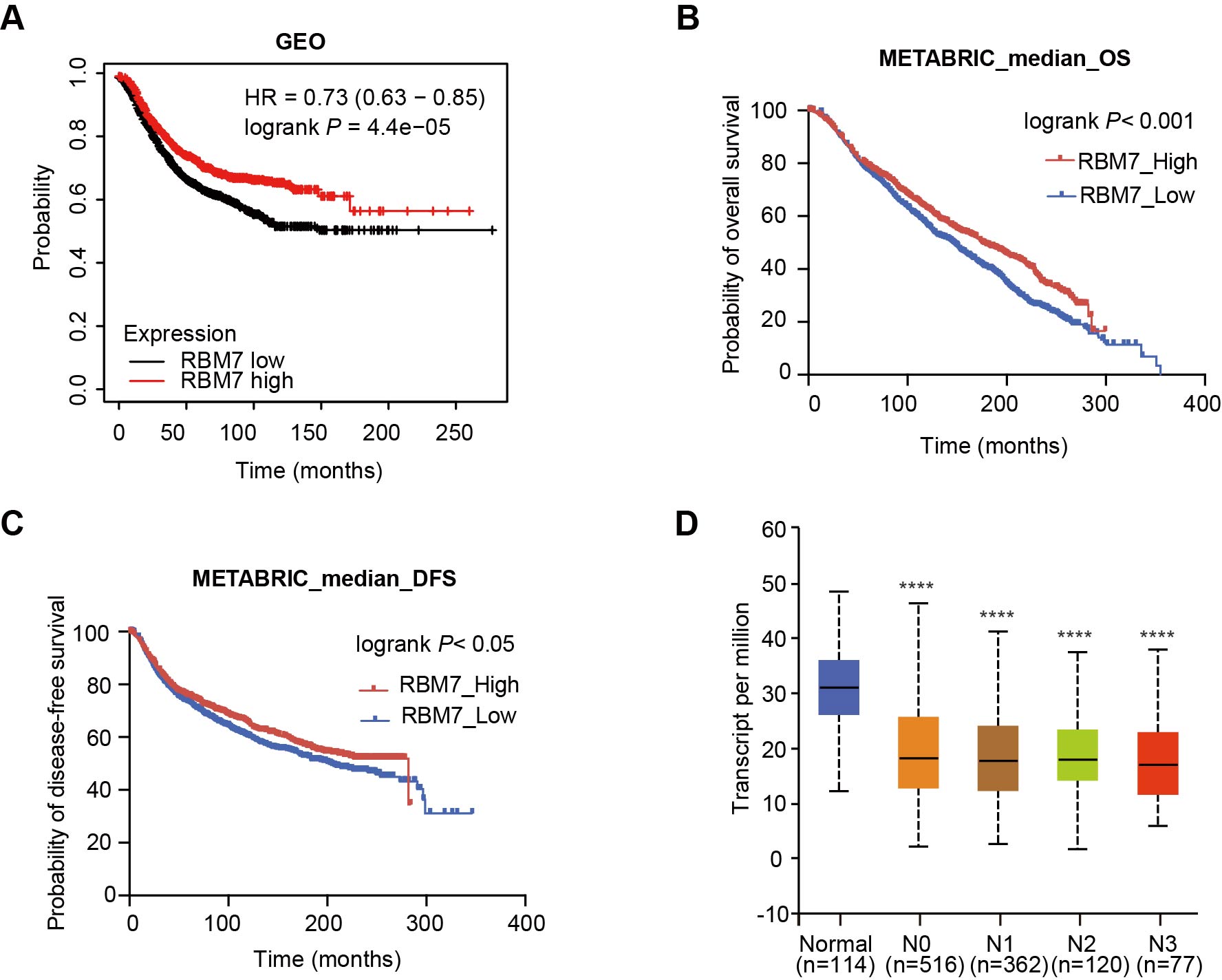

### Figure 2-figure supplement 1

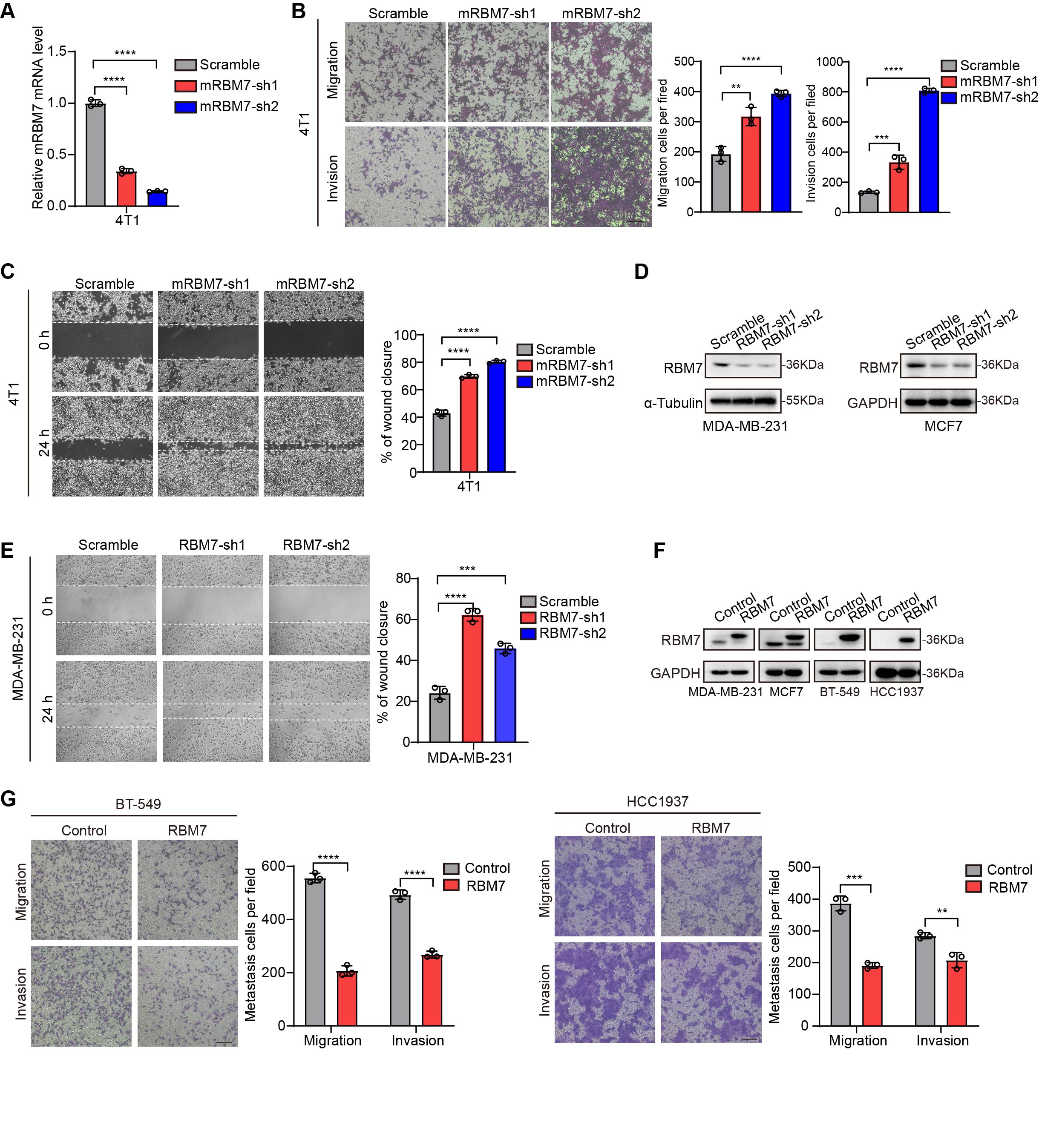

### Figure 3-figure supplement 1

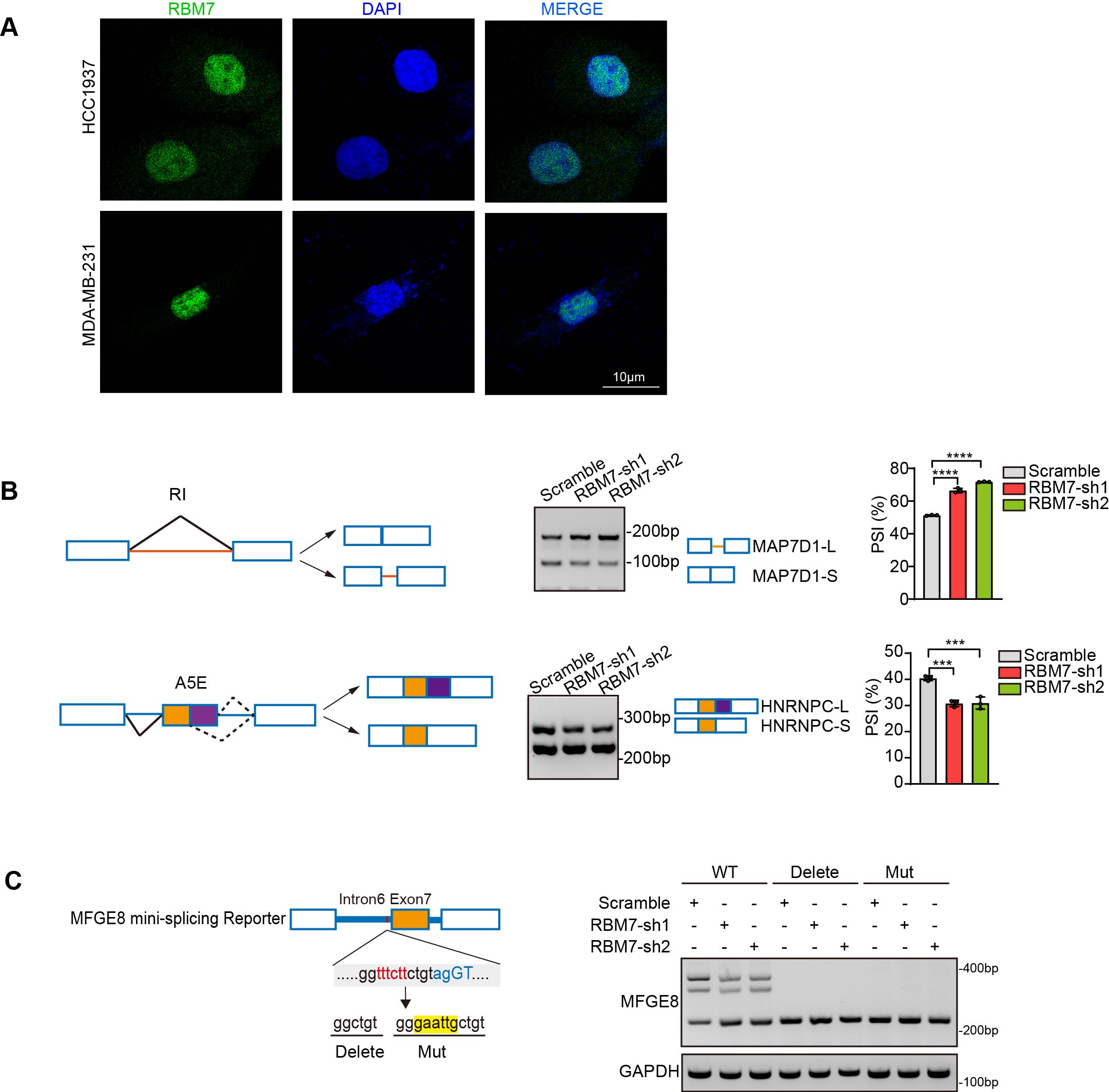

### Figure 5-figure supplement 1

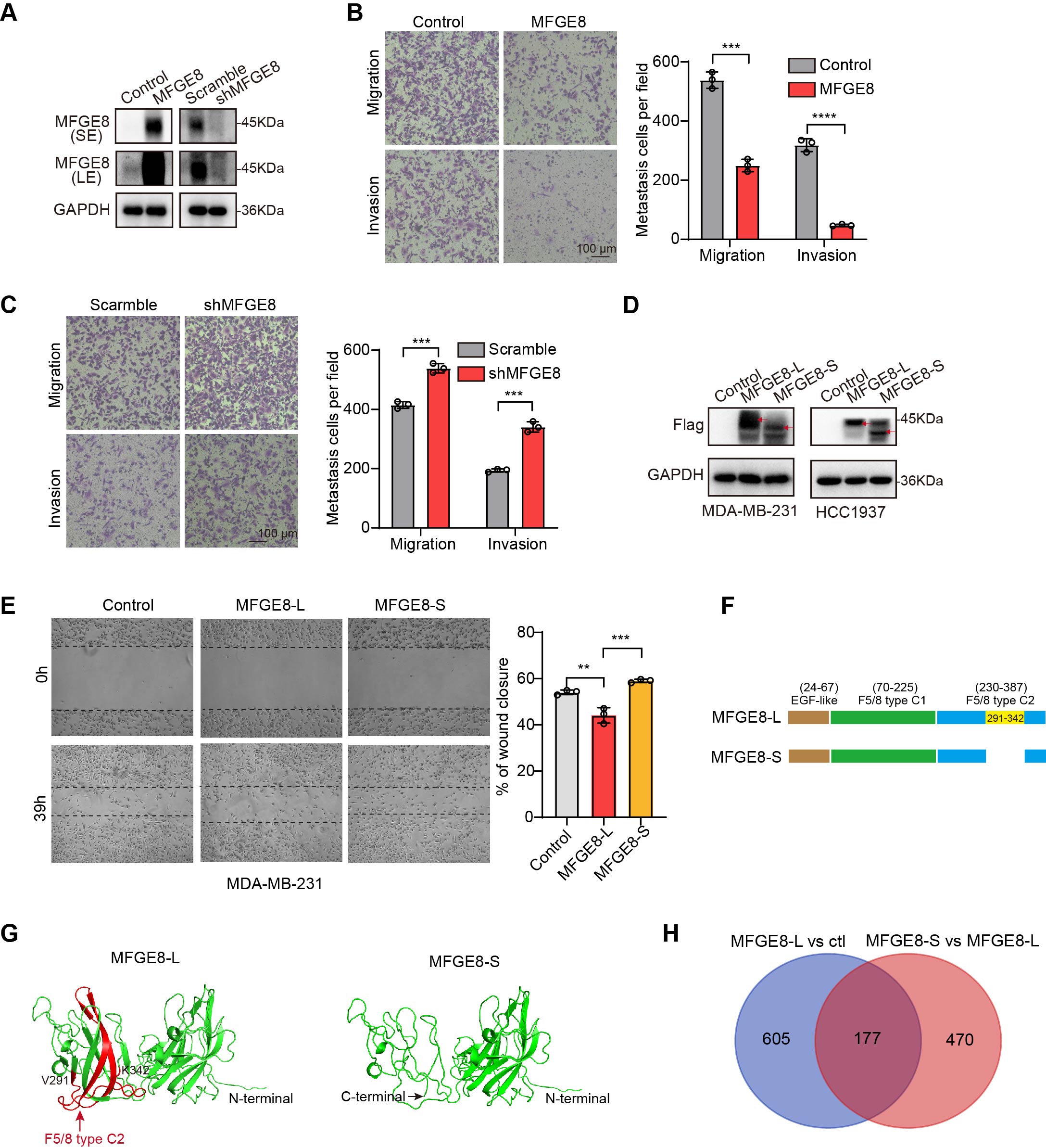

### Figure 6-figure supplement 1

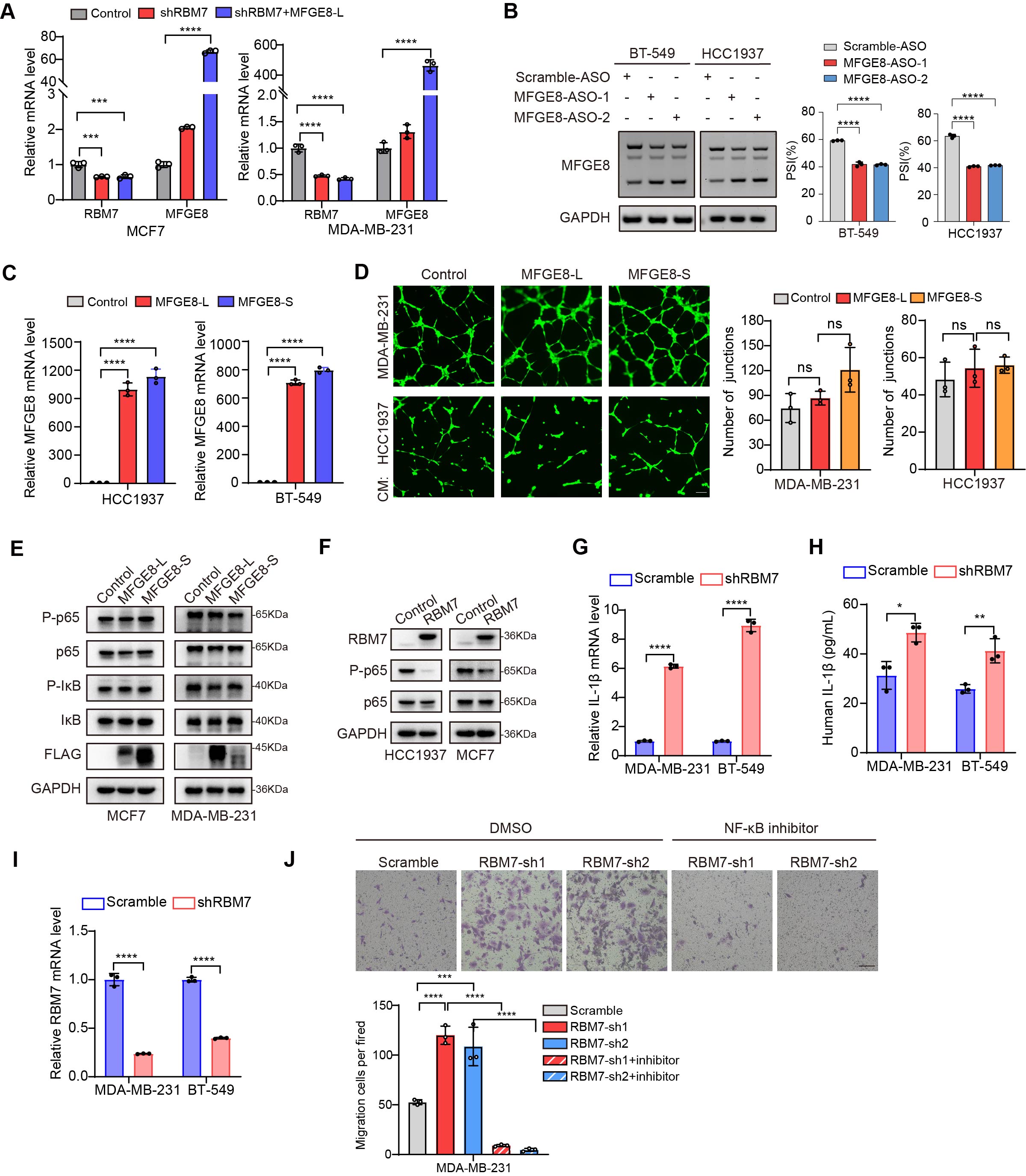

### Figure 7-figure supplement 1

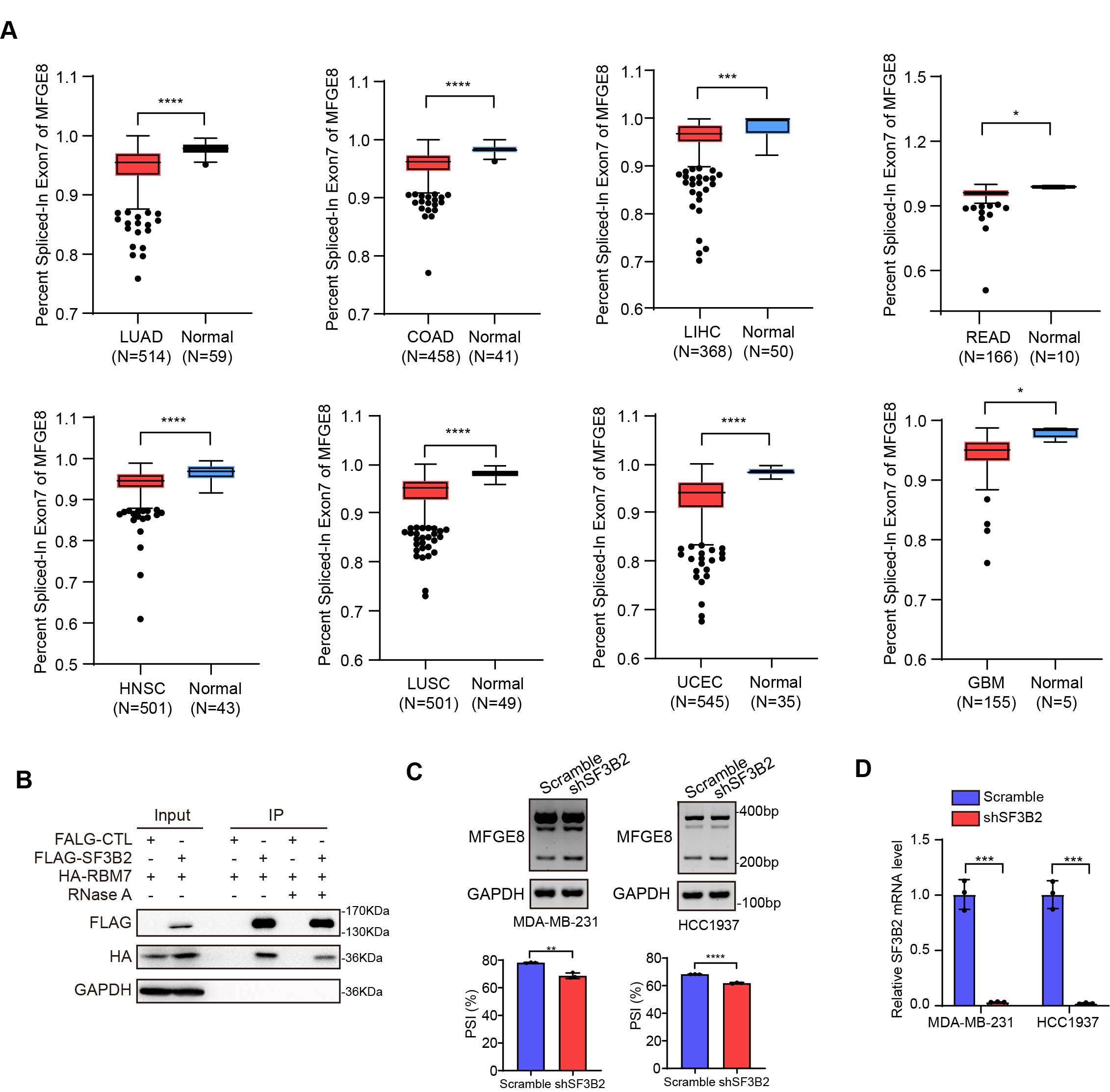
