## Supplementary Table for "RBM7 deficiency promotes breast cancer metastasis by coordinating MFGE8 splicing switch and NF-kB pathway"

| Primers | Gene | primer name | sequence |
|---|---|---|---|
| Primers for qRT-PCR | RBM7 | RBM7-qF | 5'-AAGCGGATCGCACTCTCTTT-3' |
| | | RBM7-qR | 5'-ACGCAAACTGCTTTGGTTTAC-3' |
| | GAPDH | GAPDH-qF | 5'-ATGGGGAAGGTGAAGGTCG-3' |
| | | GAPDH-qR | 5'-GGGGTCATTGATGGCAACAATA-3' |
| | CCL28 | CCL28-qF | 5'-AGAGCTGATGGGGATTGTG-3' |
| | | CCL28-qR | 5'-GGTGTTTCTTCCTGTGGCA-3' |
| | BMP2 | BMP2-qF | 5'-TCGCAGGCACTCAGGTCAG-3' |
| | | BMP2-qR | 5'-TTCCCACTCGTTTCTGGTAGTT-3' |
| | IL-1A | IL-1A-qF | 5'-GGTTGAGTTTAAGCCAATCCA-3' |
| | | IL-1A-qR | 5'-TGCTGACCTAGGCTTGATGA-3' |
| | IL-1B | IL-1B-qF | 5'-GGGCCTCAAGGAAAAGAATC-3' |
| | | IL-1B-qR | 5'-TTCTGCTTGAGAGGTGCTGA-3' |
| | LIF | LIF-qF | 5'-CGGGACCAGAAGATCCTCAA-3' |
| | | LIF-qR | 5'-ACAGCCCAGCTTCTTCTTCT-3' |
| | EBI3 | EBI3-qF | 5'-TCATCAAGCCCGACCCTCC-3' |
| | | EBI3-qR | 5'-CCAGTCACTCAGTTCCCCG-3' |
| | MMP9 | MMP9-qF | 5'-AACTTTGACAGCGACAAGAAGT-3' |
| | | MMP9-qR | 5'-ATTCACGTCGTCCTTATGCAAG-3' |
| | ITGB3 | ITGB3-qF | 5'-GGGGTAGGTTGGGAGAATGT-3' |
| | | ITGB3-qR | 5'-TCTGGGACAAAGGCTAAGGA-3' |
| | ZEB2 | ZEB2-qF | 5'-GCTGTTTCTTCGCTTCCACCT-3' |
| | | ZEB2-qR | 5'-ATTGATAAGAGCGGATCAGATGG-3' |
| | EGFR | EGFR-qF | 5'-CTGGAGAAAGGAGAACGCC-3' |
| | | EGFR-qR | 5'-TTCATCCCCCTGAATGACA-3' |
| | MMP24 | MMP24-qF | 5'-ATCTGCTTCCCTATGACTCACG-3' |
| | | MMP24-qR | 5'-CGCTTGTTTCTCCGCCTAC-3' |
| | IL-6 | IL-6-qF | 5'-GAAAGCAGCAAAGAGGCAC-3' |
| | | IL-6-qR | 5'-GCTCTGGCTTGTTCCTCACTAC-3' |
| | IL-8 | IL-8-qF | 5'-GTGCAGTTTTGCCAAGGAGT-3' |
| | | IL-8-qR | 5'-CTCTGCACCCAGTTTTCCTT-3' |
| | mGAPDH | mGAPDH-qF | 5'-CCTTCCGTGTTCCTACCCC-3' |
| | | mGAPDH-qR | 5'-GCCCTCAGATGCCTGCTTC-3' |
| | mRBM7 | mRBM7-qF | 5'-GTGTCTGTTCCCTATGCC-3' |
| | | mRBM7-qR | 5'-CACGTTACCCACTGTCCT-3' |
| | MFGE8 | MFGE8-qF | 5'-TTCCCAAGAAGTGCGAGGAGATGT-3' |
| | | MFGE8-qR | 5'-ATGCTGCAAACCCAAGAAGGTCAC-3' |
| | ZEB1 | ZEB1-qF | 5'-ATGGCGGATGGCCCCAGGTGTAAGC-3' |
| | | ZEB1-qR | 5'-GCTTACACCTGGGGCCATCCGCCAT-3' |
| | TGFB1 | TGFB1-qF | 5'-AGTTGTGCGGCAGTGGTTGAG-3' |
| | | TGFB1-qR | 5'-GGCCATGAGAAGCAGGAAAGG-3' |
| | SF3B2 | SF3B2-qF | 5'-CCCACTCCTACAGTTTTGCC-3' |
| | | SF3B2-qR | 5'-TCCAAAGCCTGGGGGATCTT-3' |
| Primers for alternative splicing | MFGE8 | MFGE8-SE-F | 5'-TGGGCCTGAA GAATAACA-3' |
| | | MFGE8-SE-R | 5'-CAAGTTCTTCTTGTGGGA-3' |
| | AKT2 | AKT2-SE-F | 5'-GCGTGTCTTCACAGAGGAGC-3' |
| | | AKT2-SE-F | 5'-AGCCCAGCAAGCAGGGACT-3' |
| | ZMYND8 | ZMYND8-SE-F | 5'-ACCCTTGACCTTTCTGGCT-3' |
| | | ZMYND8-SE-R | 5'-CCAGCCGAGACTCTTTCG-3' |
| | MBNL1 | MBNL1-SE-F | 5'-TACCAACGTG GCAATTGC-3' |
| | | MBNL1-SE-R | 5'-TAATGGGGGAAGTACAGCTT-3' |
| | MAP7D1 | MAP7D1-RI-F | 5'-ATCGCAGCCTGCAGCTGA-3' |
| | | MAP7D1-RI-R | 5'-TGCACGCTTCGGGTGACGCT-3' |
| | HNRNPC | HNRNPC-A5E-F | 5'-GGCTTTGCCTTCGTTCAG-3' |
| | | HNRNPC-A5E-R | 5'-CATCCTATCATAATAGTCCCGTTG-3' |
